## Supplementary figures and images for "The proteomic landscape of resting and activated CD4+ T cells reveal insights into cell differentiation and function"

### Supplementary Figure 1

#### Supplementary figure 1

A.

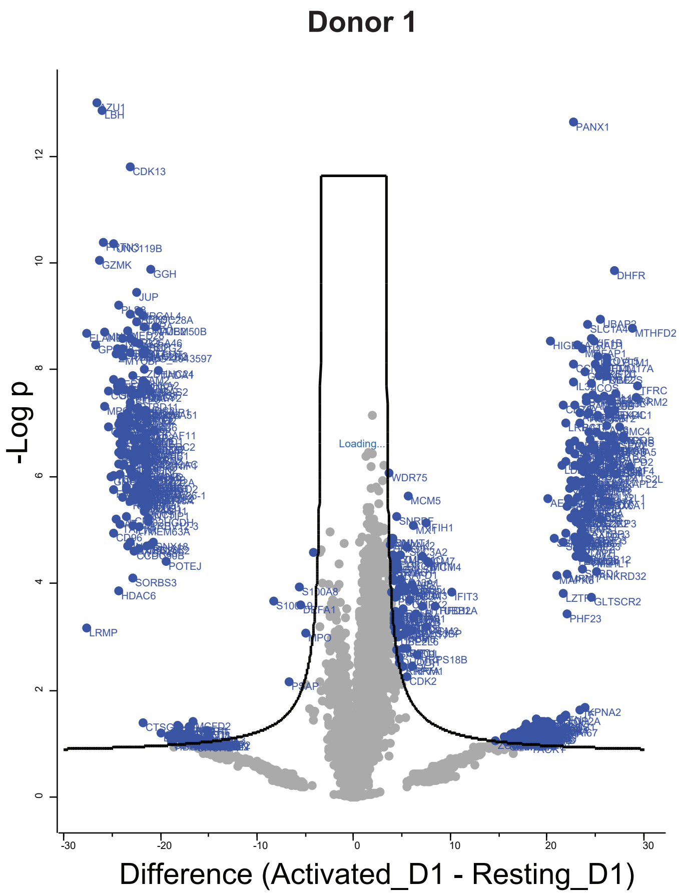


B.

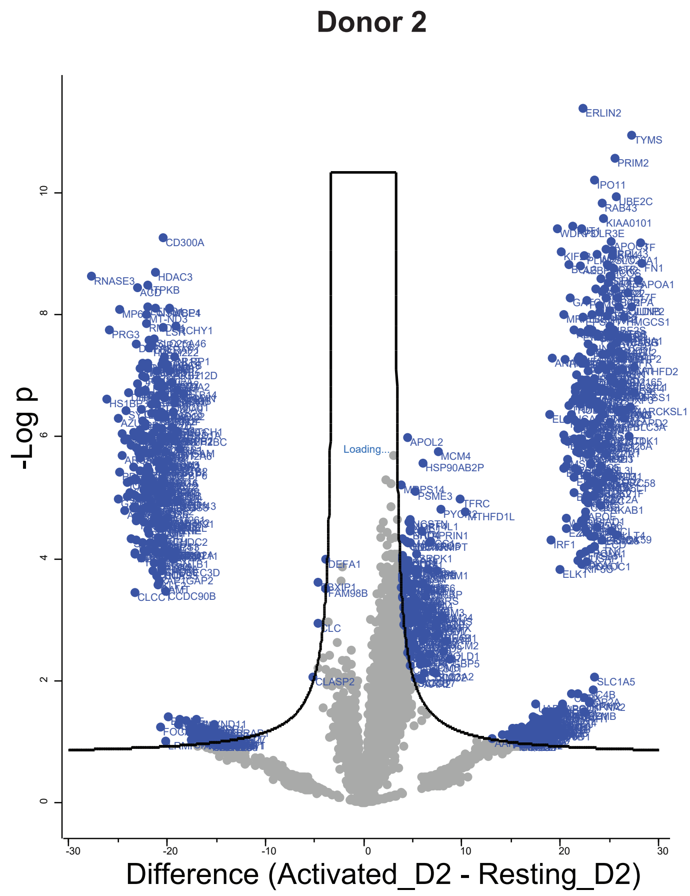

### Supplementary Figure 2

**Supplementary figure 2**

**A.** **Downregulated proteins in Donor 1**

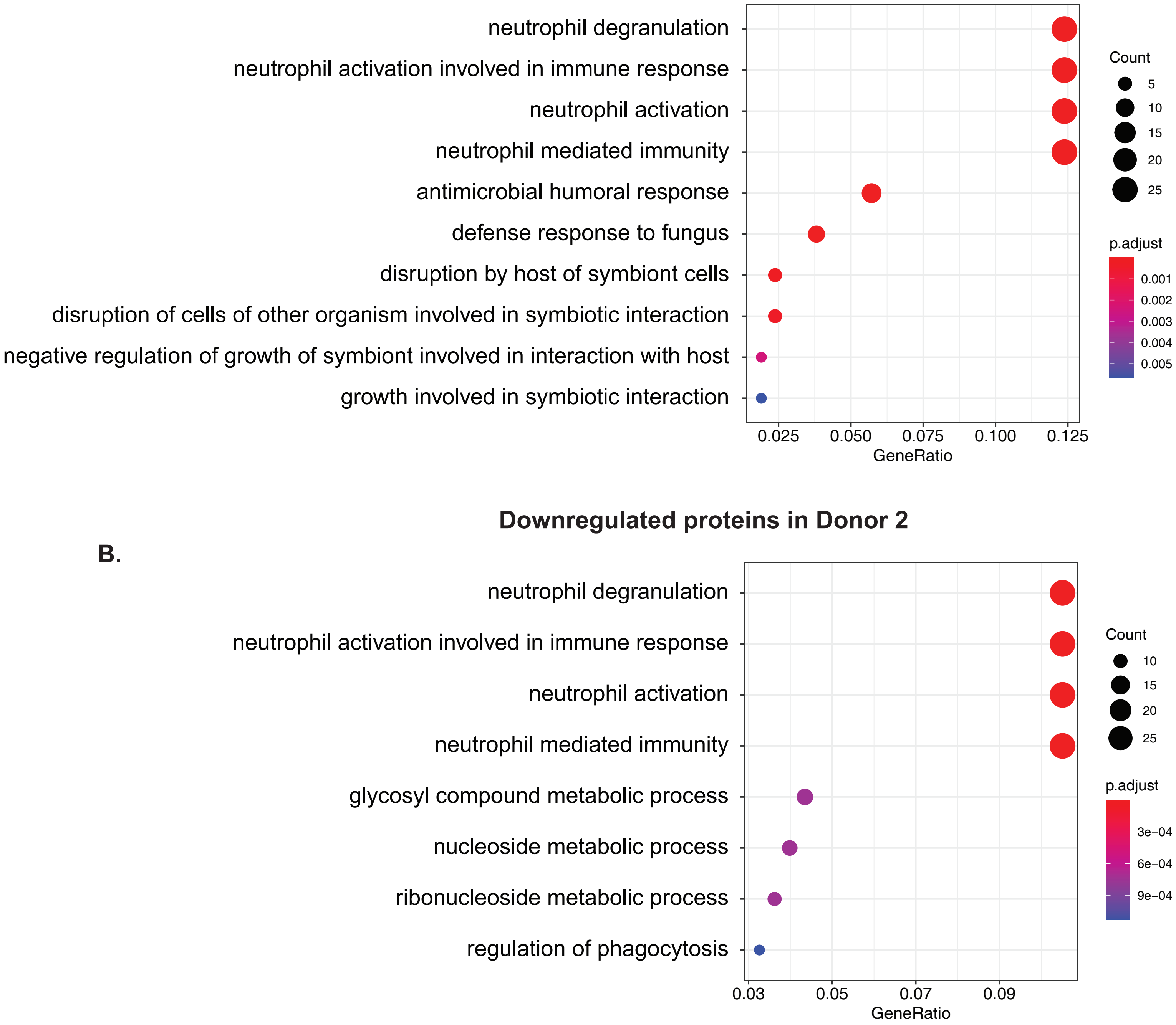


**Downregulated proteins in Donor 2**

**B.**

### Supplementary Figure 3

Supplementary figure 3

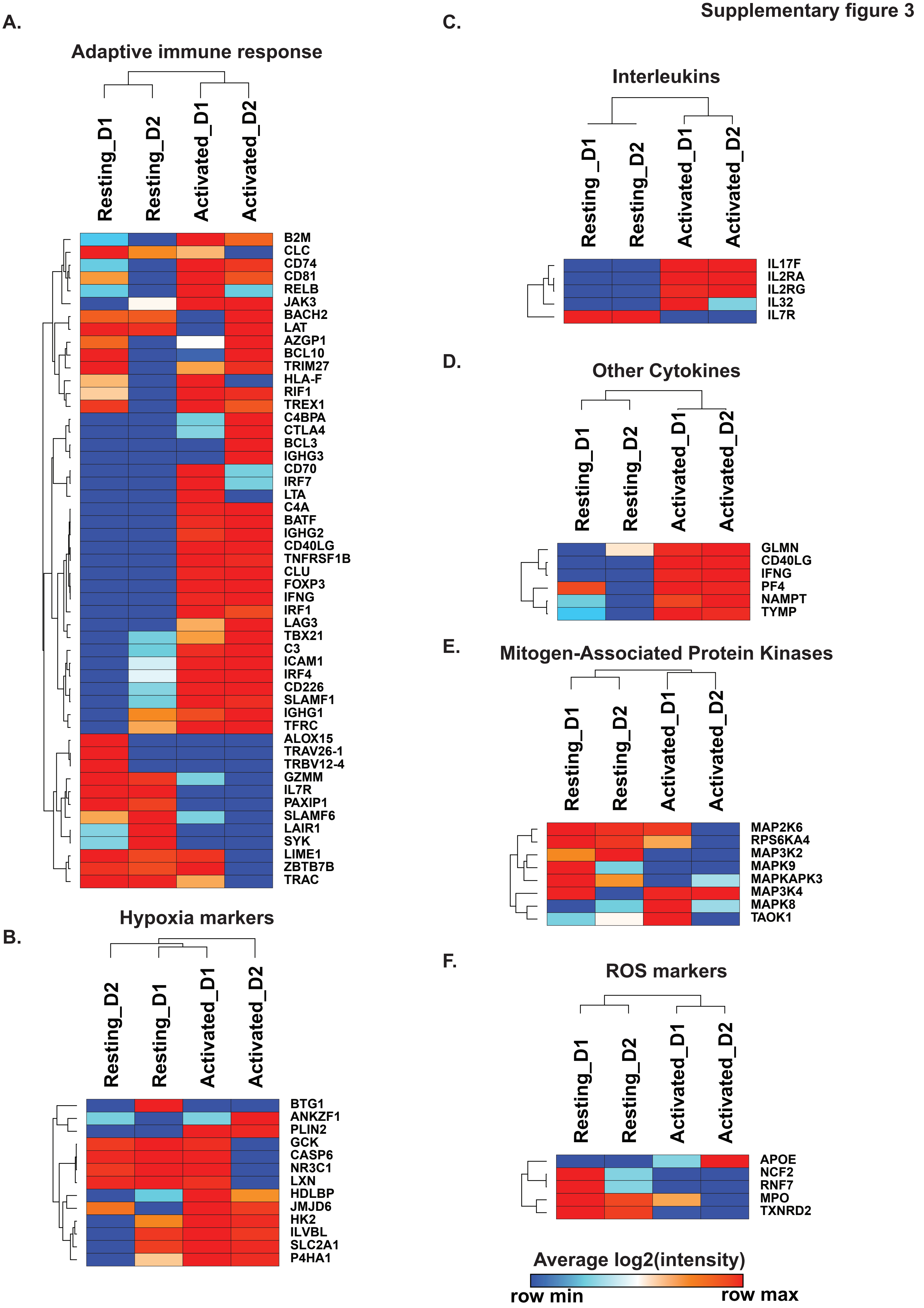

### Supplementary Figure 4

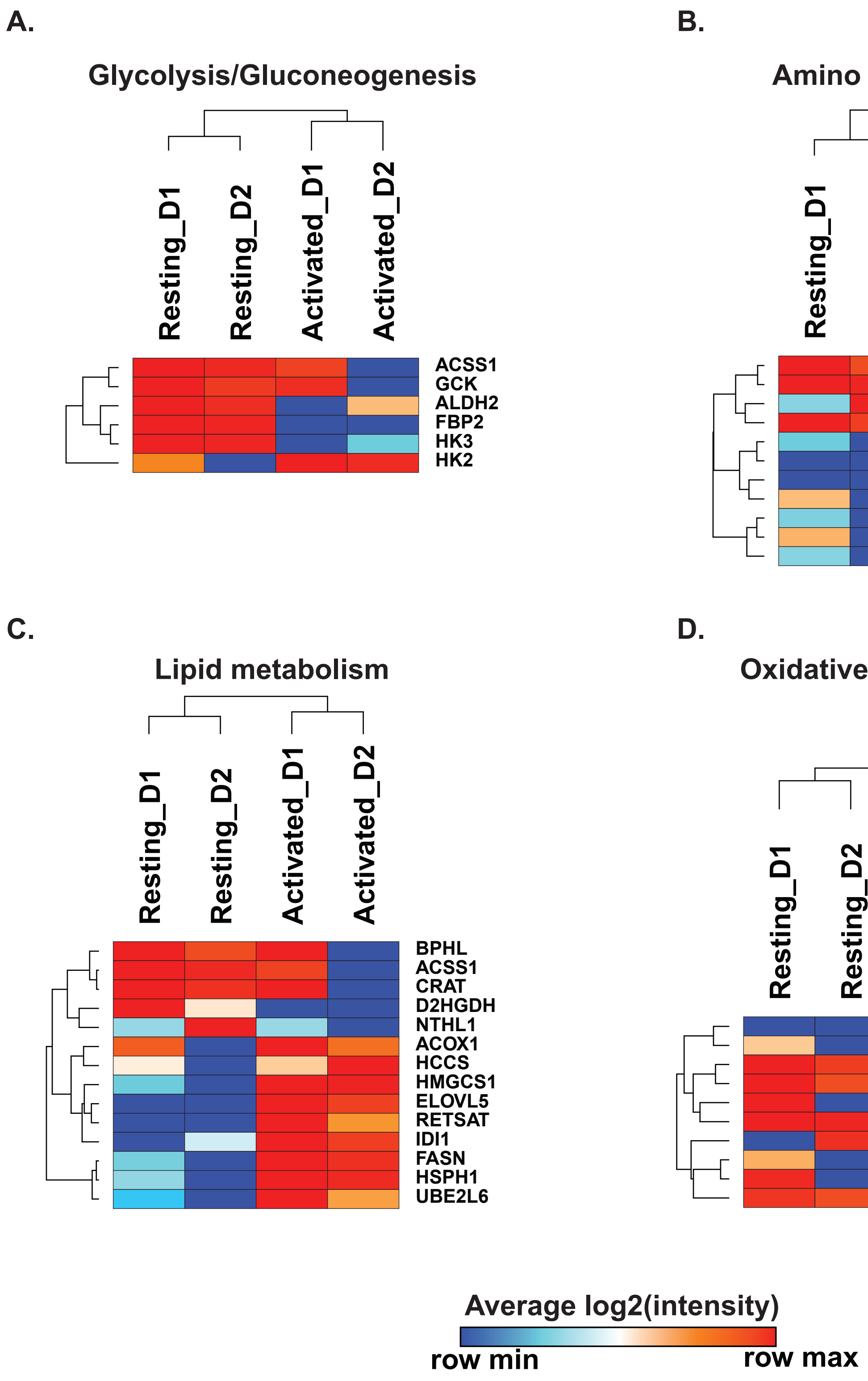

### Supplementary Figure 5

Supplementary figure 5

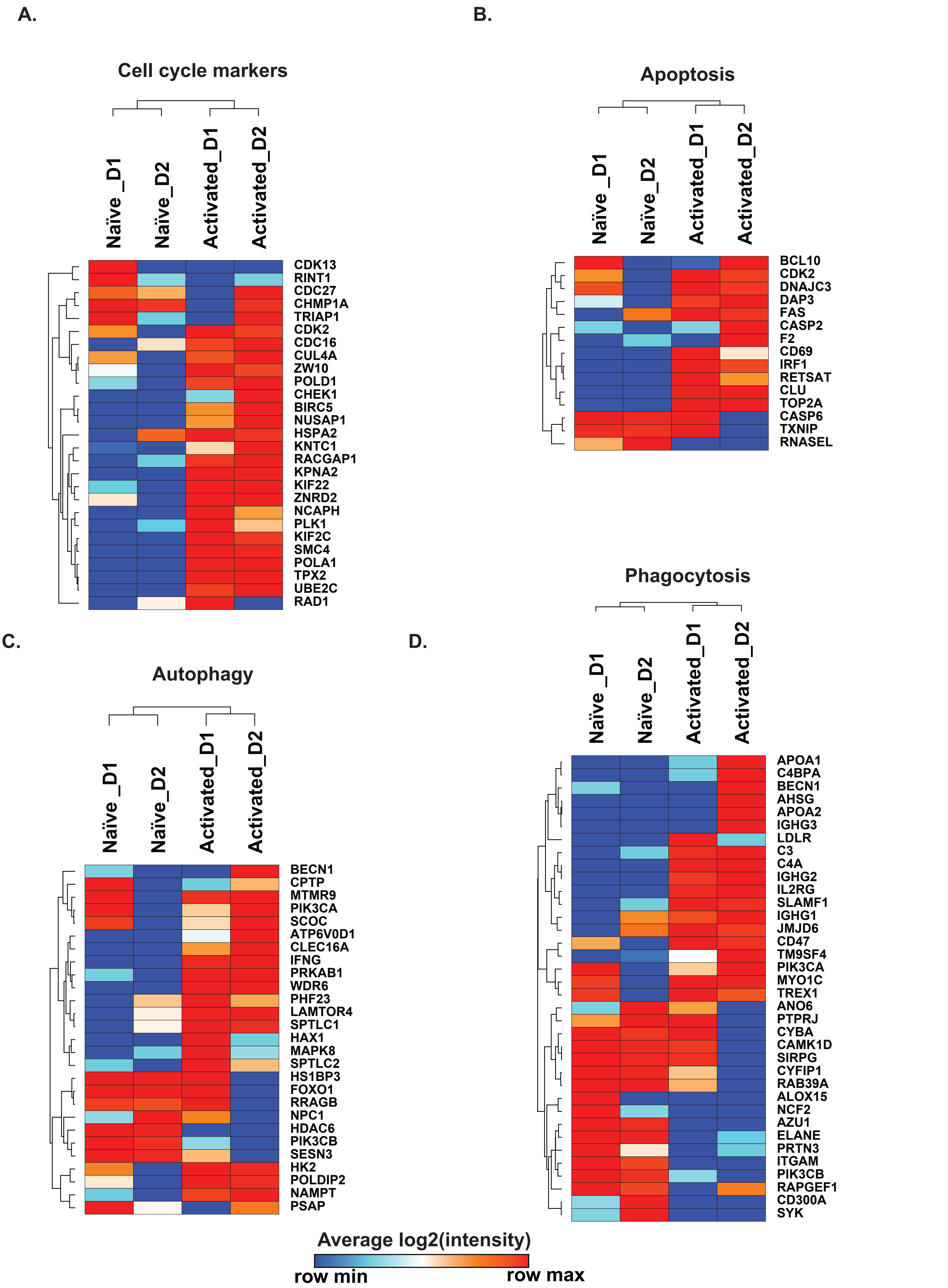
